# A cost-efficient open source laser engine for microscopy

**DOI:** 10.1101/796482

**Authors:** Daniel Schröder, Joran Deschamps, Anindita Dasgupta, Ulf Matti, Jonas Ries

**Author notes:** These authors contributed equally. https://github.com/ries-lab/LaserEngine (Daniel Schröder, Joran Deschamps).

## Abstract

Scientific-grade lasers are costly components of modern microscopes. For high-power applications, such as single-molecule localization microscopy, their price can become prohibitive. Here, we present an open-source high-power laser engine that can be built for a fraction of the cost. It uses affordable, yet powerful laser diodes at wavelengths of 405 nm, 488 nm and 640 nm and optionally a 561 nm diode-pumped solid-state laser. The light is delivered to the microscope via an agitated multimode fiber in order to suppress speckles. We provide the part lists, CAD files and detailed descriptions, allowing any research group to build their own laser engine.

## 1. Introduction

Fluorescence microscopy is a central method in biology and modern implementations often use lasers as illumination sources. In order to image different fluorescent labels, several laser lines spanning the visible range are required. For stability purposes, the different laser beams are often overlaid within a laser combiner, whether commercially sold or custom-built, and conveyed to the microscope by an optical fiber. Because of their cost, scientific-grade lasers used in advanced microscopes often account for a considerable portion of the total microscope price [1], with single units reaching thousands to tens of thousands of euros/dollars. This is especially the case in power-hungry superresolution methods such as single-molecule localization microscopy (SMLM) [2–4], which typically requires a power density in the range 1-100 kW/cm^2^[5].

As a substitute for scientific-grade lasers, inexpensive laser diodes can deliver comparable laser power at a fraction of the cost (<100 euros/dollars). Such laser diodes have already been successfully used for low cost illumination in photoacoustic microscopy [6–9], pump-probe microscopy [10], stimulated emission-depletion (STED) microscopy [11] and SMLM [12]. Due to their high divergence and asymmetrical intensity profile, they are challenging to couple into single-mode optical fibers with a decent efficiency. They are therefore often coupled into multimode fibers, leading to higher coupling efficiencies and a more uniform profile. Homogenous illumination is crucial in SMLM [13, 14], as it leads to constant photophysics of single molecules and thus constant image quality across the field of view. However, a consequence of coupling laser light into multimode optical fibers is the appearance of laser speckles [15]: small-scale heterogeneities in the illumination that degrade the illumination profile. Perturbing the speckle pattern faster than the camera frame rate effectively averages it out. Several approaches have been explored in order to remove speckles from a multimode fiber profile, including rotating optical elements [12, 16], oscillating diffusive membranes [14, 17] or mechanical agitation [12, 18–20].

In this work, we developed an affordable high-power laser engine using inexpensive laser diodes, a multimode fiber and speckle-reduction methods. The laser engine is compatible with any wide-field microscope, commercial or custom-built. Four laser diodes (405 nm, 488 nm and 2x 638 nm) deliver from 25 mW (405 nm) to 1 W (638 nm) of output power per line. Since inexpensive laser diodes for red fluorophores or fluorescent proteins are not available, we added a powerful diode-pumped solid-state laser (dpss, 561 nm, 500 mW). Building on well-established methods for homogenization of multimode fibers, we assembled a simple fiber agitation module for our laser engine. We show that this system efficiently suppresses laser speckles and that the high laser power allows performing high-speed SMLM on mammalian cells.

Most of the components are stock parts, while widely available alternatives exist for the custom elements. The parts list and all designs are freely available online (Github, https://github.com/ries-lab/LaserEngine), enabling other researchers to easily assemble their own laser engine for a fraction of the cost of a commercial instrument.

## 2. Results

### 2.1. Laser engine

Fig. 1 shows a schematic of the laser engine beam path (see also Appendix Fig. 1 (a) for pictures). Unless otherwise stated, all parts were purchased from Thorlabs. The laser engine is built on a 300 mm by 450 mm aluminum breadboard (MB3045/M). The laser diodes are mounted on a custom aluminum holder (see Appendix Fig. 1 (b)) with thermal paste and held by passive heat sinks. The holder allows mounting up to five diodes in parallel. We opted for two 638 nm diodes (HL63193MG, Oclaro/Ushio), each delivering 700 mW, in order to cover high power applications [13, 18, 21]. In addition to the red channel, we added a UV (405 nm, 80 mW, dl-7146-301s, SANYO) and a 488 nm (55 mW, BLD-488-55, Lasertack) laser diode. Within the holder, 8 mm aspherical lenses (A240TM-A) are used to collimate the output of the laser diodes (see arrow on Appendix Fig. 1 (b)). The two red beams are set to propagate with perpendicular polarizations by adjusting the orientation of the diodes (see polarization in Fig. 1). They are then combined using a polarizing beam splitter (PBS251). Besides the laser diodes, we added a high power 561 nm dpss laser (500 mW, gem561, Laser Quantum). All laser lines are then overlaid using dichroic mirrors (DMLP605, DMLP505 and DMLP425). The beams are finally coupled into a square (150×150 μm, 0.39 NA, M103L05) or round (105 μm, 0.22 NA, FG105UCA custom cable with FT038 tubing) multimode optical fiber using a 19 mm lens (AC127-019-A). To facilitate alignment and coupling into the fiber, each optical element is centered on a breadboard hole and held either by a precision (mirrors, KS05/M or KS1) or a flexure (dichroic mirrors, PFM1/M) mount. This compact design circumvents the need for cylindrical lenses. We obtained coupling efficiencies in the square multimode fiber of 47% at 405 nm (which leads to an estimated 12 mW at the sample given the measured loss before the objective and the objective transmittance at this wavelength), 80% at 488 nm (24 mW), 81% at 561 nm (320 mW) and 80% at 638 nm (2× 370 mW).

**Fig. 1.**
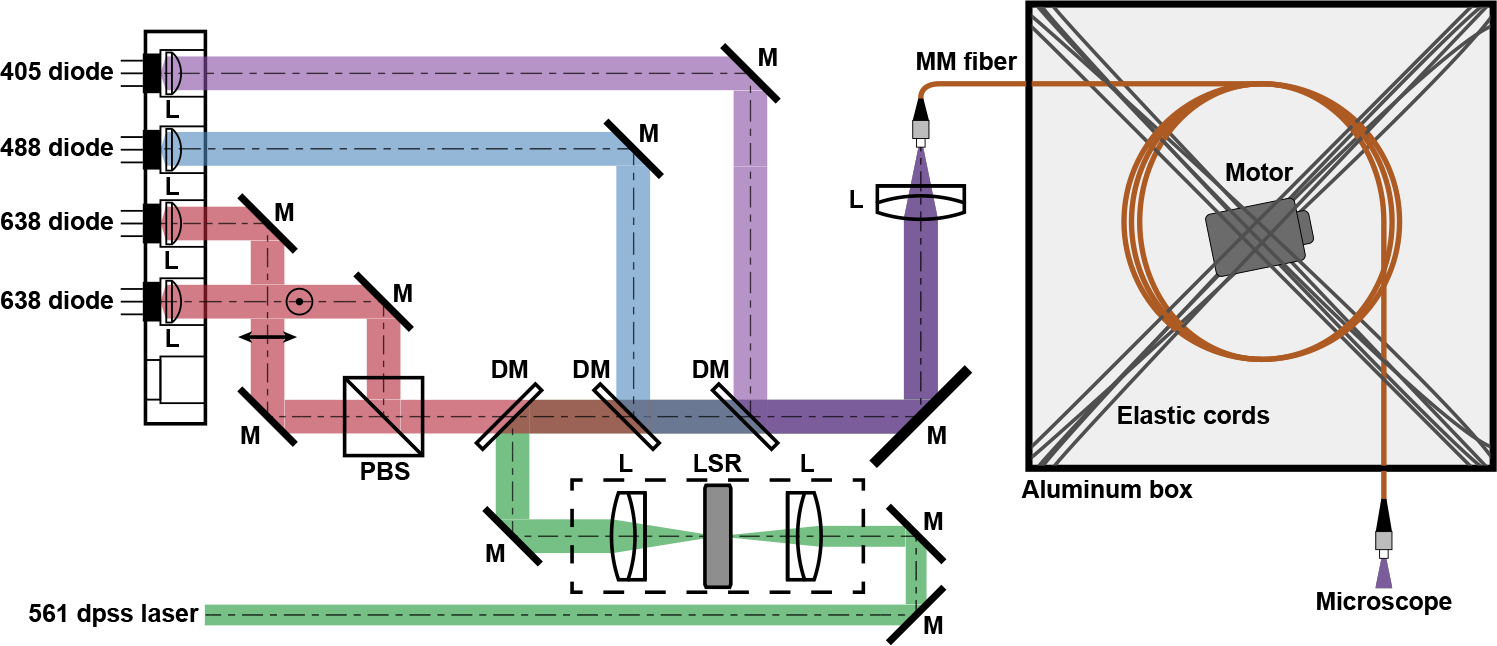
Optical path of the laser engine. The laser diodes are mounted in a custom holder. They are collimated by an aspherical lens and combined using a polarizing beam splitter (red diodes) and dichroic mirrors. The diode-pumped solid-state (dpss) laser is focused on a speckle-reducer and then collimated by achromatic lenses. All lasers are coupled into a multimode optical fiber. The fiber is then introduced into a heavy aluminum box and intertwined with elastic cords suspended on the box corners. A vibration motor is held at the center of the box by the same elastics. Finally, the output of the optical fiber is used in the illumination path of the microscope. The arrow and circle on the red diodes’ outputs denote the p and s polarizations. L: lens, M: mirror, PBS: polarization beam splitter, DM: dichroic mirror, LSR: laser speckle-reducer, MM: multimode. The dashed rectangle denotes the optional speckle-reduction system for the dpss laser.

A laser diode driver (eu-38-ttl, Roithner) controls each laser diode using a single input channel (transistor-transistor logic (TTL) signal). The output power of the diodes can be adjusted using a potentiometer on the board. In order to control the power of the diodes from the computer, we implemented an additional electronic board to convert the TTL signal and a pulse-width modulation (PWM) signal coming from a field-programmable gate array (FPGA, Mojo, Alchitry), or an Arduino board, to an analog signal. We modified the laser diode driver to bypass the potentiometer by directly feeding in the analog signal to components on the board (see the Github repository for in-depth details), while limiting the maximum current supplied to the laser diode in order to prevent damages. Using these electronics, we could pulse the diodes in the hundred of microseconds range.

### 2.2. Coupling into the microscope

To achieve homogeneous illumination, the output of the multimode fiber is imaged onto the sample plane [14]. To do so, the laser light must be collimated before entering the objective. Most commercial wide-field or total-internal fluorescence (TIRF) instruments feature a port to access the illumination path of the microscope. If a lens is already present in the illumination path before the objective, then the output of the optical fiber must be focused in the object plane of the lens. In our custom microscope, we used a single lens in front of the optical fiber to form its output image in the focal plane of an illumination lens; the latter being aligned in a 4f configuration with the objective. The size of the illuminated field of view is adjusted by changing the optical path length between fiber exit and illumination lens, then by fine-tuning the position of the first lens to refocus the fiber output in the sample. Two mirrors are used in between to move the illumination laterally. We set the illuminated area in the sample to about 32×32 μm^2^. Since we observed auto-fluorescence in the multimode optical fiber, we placed a four-line clean-up filter (390/482/563/640 HC Quad, AHF) before the illumination lens.

### 2.3. Speckle-reduction

The propagation of multiple coherent laser modes in the multimode fiber results in speckles in the output beam profile. Speckles are more pronounced for lasers with longer coherence length. It is possible to average speckles out and to achieve an effective homogeneous illumination by physically moving the optical fiber at high frequencies. To this end, we suspended a vibrational motor (DC 1.5-6V, 16.500 rpm, 22g, Sourcingmap) with elastic cords (19710640, Prym) in an aluminum box (see simplified depiction in Fig. 1 and picture in Appendix Fig. 1 (d)). The optical fiber was then intertwined with the elastics. As a result, turning the motor on produces strong vibrations in the elastic cords and agitates the optical fiber in a complex manner. The aluminum enclosure dampens the motor noise to comfortable levels for the user. In order to visualize in detail the effect of the speckle-reduction, we imaged the output of the multimode fiber directly onto a CMOS camera (DCC1545M) using a single achromatic lens (30 mm, AC254-030-A). Fig. 2 (a) shows the speckle pattern obtained with each laser line using the square multimode fiber. The 638 nm diodes show speckles of low magnitude while the 405 nm and 488 nm, on the other hand, display stronger speckle patterns. Dpss lasers are notorious for their longer coherence length and consequently the 561 nm output presents speckles of larger amplitudes than the other lasers. Without speckle reduction, the speckle contrast (indicated at the bottom of each image on Fig. 2 (a), see methods for the definition) of the 561 nm laser is 0.22, higher than that of the 405 nm and 488 nm diodes (0.14 and 0.13, respectively) and of the 638 nm diodes (0.05 for each). With fiber agitation and 5 ms exposure time, speckles are averaged out for all laser lines (see Fig. 2 (b)), with small residual speckles only observed for the 561 nm laser. The speckle contrasts indeed substantially drop to 0.020 (405 nm), 0.022 (488 nm), 0.031 (561 nm) and 0.020 (for both 638 nm diodes). Using both 638 nm laser diodes simultaneously further reduces the speckle contrast of the illumination. The lateral profile of the fiber outputs with and without agitation allows comparing the amplitude of the speckles to the large-scale inhomogeneity of the illumination (see Fig 2 (c)). Both 405 nm and 638 nm diodes display a flat profile while the other laser lines show a stronger intensity drop at the edges. The effect of the speckle-reduction can be appreciated by comparing intensity profiles with and without agitation. Note that, for all lasers, the corners of the square fiber output are of slightly lower intensity.

**Fig. 2.**
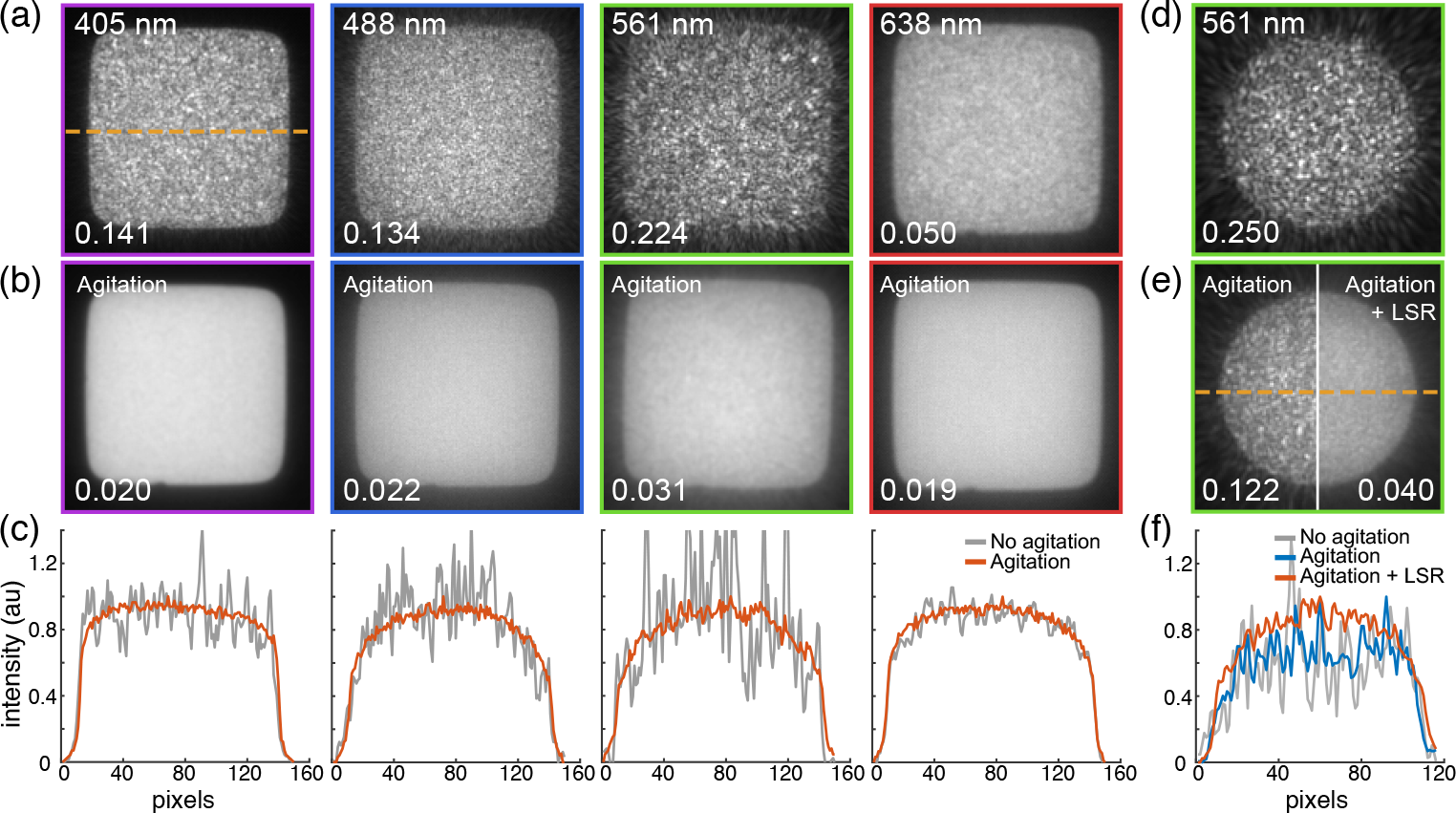
Speckle-reduction. (a) Profiles of the square multimode fiber output without speckle reduction for the four laser lines (only one 638 nm diode is shown here). The speckle contrast is indicated at the bottom of each image. (b) Same profiles with agitation. (c) Lateral intensity profiles of (a) (grey curve) and (b) (orange curve) along the dashed line of (a), averaged over a width of 4 pixels. (d) Profile of the round multimode fiber output for the 561 nm dpss laser without speckle-reduction. (e) Same with agitation (left) and with both agitation and the LSR on (right). (f) Lateral line profiles of (d) (grey curve) and of the bottom panels of (e) (left and right corresponding to the blue and orange curves respectively) along the dashed line of (e), averaged over a width of 4 pixels. All images were acquired with 5 ms exposure time.

The amplitude of the speckles, and therefore the measured speckle contrast, depends on the output power of the laser diodes (see Appendix Fig. 2). In particular, we observed that at their maximum power, the laser diodes display speckles of smaller amplitude than at low power. At 5 ms exposure time, the speckles are suppressed with similar efficiency regardless of the output power. Furthermore, the speckles are efficiently suppressed at exposures as low as 0.1 ms (see Appendix Fig. 3) for all laser diodes. In the case of the dpss laser, the speckles remain visible below 1 ms exposure time and the efficiency of the agitation rapidly decreases.

We additionally tested the system with a round multimode fiber (105 μm, 0,22 NA, FG105UCA with FT038 tubing). Overall, the fiber outputs had more pronounced speckles and bell shape profiles (see Appendix Fig. 4) than that of the square optical fiber. The agitation alone did not prove as efficient, in particular for the 561 nm laser. Fig. 2 (d) shows the dpss laser output with the round fiber without speckle reduction. There, the speckle pattern is slightly stronger than with the square fiber. Substantial speckles remain even during fiber agitation (Fig. 2 (e), left panel). In order to further break the coherence, we followed previous work [14] and added a commercial laser speckle-reducer (LSR, LSR-3005-17S-VIS, Optotune) in the beam path of the 561 nm dpss laser (see Fig. 1 (a), dashed box). As reported earlier, this decreases the coupling efficiency at the multimode fiber due to an increased divergence of the laser beam, in addition to the lower NA and smaller core of the fiber. We obtained a coupling efficiency of about 50%. The use of the LSR in conjunction with the agitation leads to a better speckle-reduction (Fig. 2 (e), right panel), with a three-fold improvement of the speckle contrast (0.12 to 0.04). The fiber output after speckle-reduction with both agitation and LSR is smoother than with agitation alone, exemplified by the intensity profiles in Fig. 2 (f). The LSR did not lead to a perceptible improvement of the speckle-reduction in the case of the square fiber.

### 2.4. Application to fast SMLM

Because SMLM relies on the accumulation of thousands of images with sparse single-emitters, obtaining a superresolved image is slow compared to other microscopy methods. Organic dyes such as AF647 can be driven to fast blinking dynamics with high laser power and fast cameras [5, 21, 22], leading to a several-fold improvement of the imaging speed. Our laser engine can deliver the laser powers required for such experiments. As a proof-of-concept, we imaged Nup96, a protein of the nuclear pore complex that was labeled with AF647 via a SNAP tag [23, 24], and acquired SMLM images in less than 40 s. The reconstructed lower nuclear envelope image is shown on Fig. 3 (a), with hundreds of nuclear pores distributed over the surface. Most of them exhibit a ring structure, highlighting the quality of the experiment. The median of the filtered localization precision (filtering out localization with a precision > 15 nm) is 3.9 nm. Fig. 3 (b) shows two close-up regions of interest (Roi 1 and 2 in Fig. 3 (a)). Both regions contain fully formed rings as well as nuclear pores with lower coverage of localizations. We performed an analysis based on automated detection of NPC sites, removal of low quality pores and least-square fitting of a circle to the nuclear pores localizations (as in [24]) within the 10 × 5 μm^2^ dashed region of Fig. 3 (a). The average diameter of Nup96 obtained is 112.7 ± 5.4 nm (N=211), in good agreement with the current literature [25] and in particular with experiments carried out with scientific-grade lasers and optimized imaging conditions [24].

**Fig. 3.**
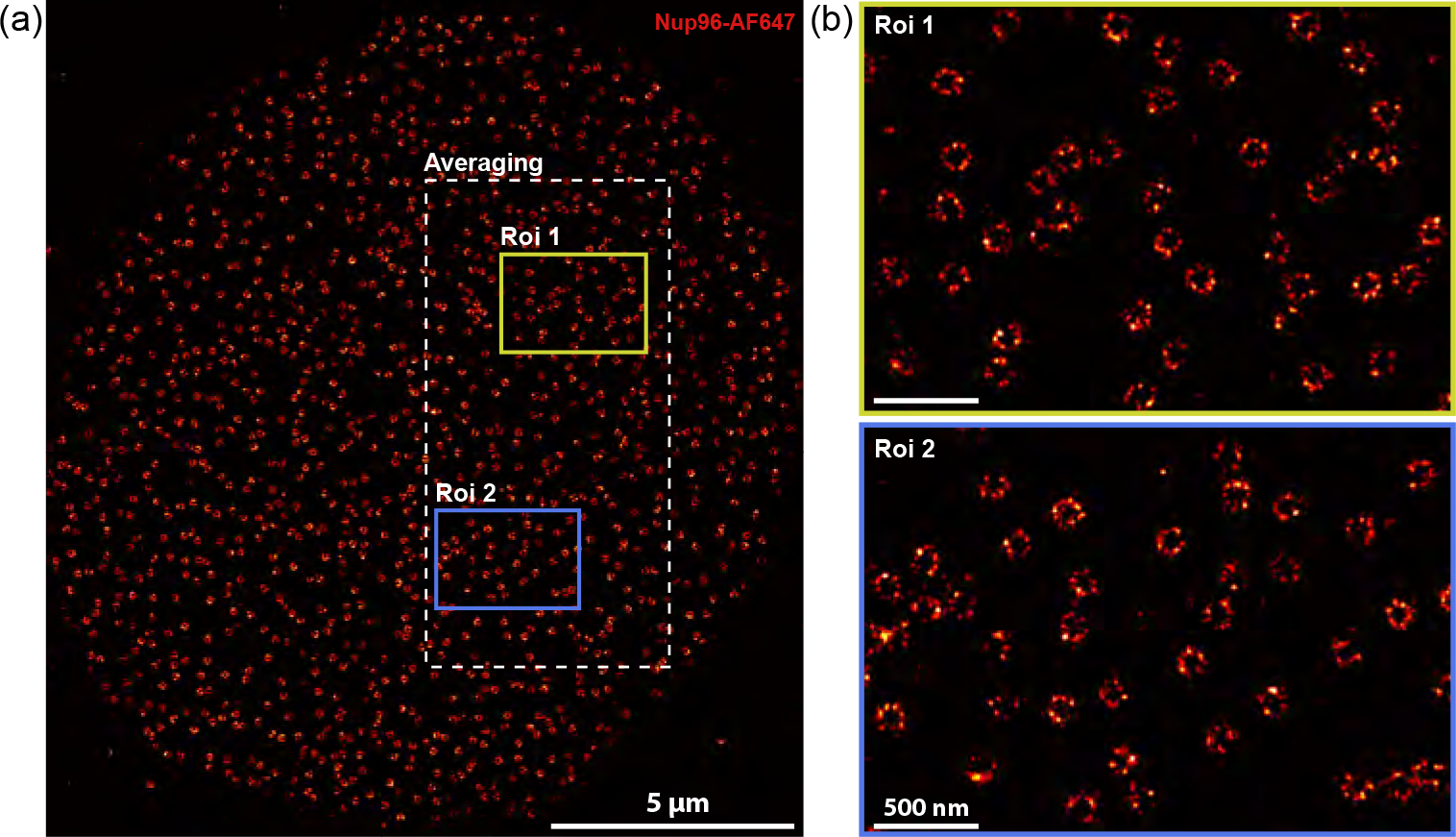
High-speed localization microscopy. (a) Superresolved image of a fixed U2OS cell nucleus, with the nucleoporin Nup96 tagged with AF647 via SNAP-tag and imaged in a dSTORM buffer (see methods) with a laser power of 30 kW/cm^2^, reconstructed from 3000 frames with an exposure time of 10 ms. The dashed rectangle delimitates the area used for averaging the diameter of individual nuclear pores. (b) Close-up images of two regions of (a): Roi 1 (yellow rectangle) and Roi 2 (blue rectangle).

## 3. Discussion

Here, we presented a four-color high-power laser engine designed to be easily assembled in any laboratory. The optical elements cost approximately 3000 euros/dollars in total (excluding the 561 nm dpss laser, the custom mechanical pieces and electronics, and the LSR). This is in stark contrast to commercial laser systems. For instance, a scientific-grade 640 nm laser diode typically costs several thousands of euros/dollars for powers up to 200 mW and tens of thousands of euros/dollars for 1W, while the 700 mW 640 nm laser diode used in this manuscript costs less than 100 euros/dollars. The laser diodes, their drivers and the vibrational motor can be purchased from a variety of distributors (Thorlabs, Roithner, Lasertack, Lasershop and Sourcingmap).

The custom laser diode holder is a complex anodized aluminum piece designed to guarantee stability, small physical footprint and efficient heat dissipation. Operating the diodes at high power without any cooling system will result in immediate shut down or even permanent damages. The conjunction of the aluminum mount and individual heat sinks not only allows high lasing power but also ensures a stable output power (see Appendix Fig. 5). The custom laser diode holder can be replaced by commercially available temperature-controlled mounts (e.g. Thorlabs, LDM56/M).

Aside from the low-cost laser diodes, we added a scientific-grade dpss 561 laser to the laser engine. Laser diodes are currently not available in the 560 nm region of the spectrum at equivalently low price. Since many fluorescent proteins used in SMLM require such an illumination wavelength (e.g.: mMaple [26], mEos3.2 [27]), this laser line is often a necessary addition. Our diode holder and beam path currently leave the possibility to add a fifth laser diode in case such a wavelength becomes available.

Furthermore, we compared two multimode fibers of different geometry, square (150 μm, 0.39 NA) and round (105 μm diameter, 0.22 NA). Because the profile of the low-cost laser diodes is asymmetrical and propagates with strong divergence, coupling efficiencies in the round optical fiber were limited to the 50% to 70% range depending on the diode. The larger core and NA of the square fiber led to coupling efficiencies around 80%, except for the 405 nm diode for which chromatic aberrations and overall transmission yielded a coupling efficiency below 50%. In addition, the square fiber had superior performances in both output shape (more top-hat than bell-curved) and speckle-reduction for all laser lines. We found that the speckle reducer (LSR) improved the homogeneity of the dpss laser when used with an agitated round multimode fiber, at the price of lower coupling efficiency. On the other hand, it only had a marginal effect on the square fiber and was therefore removed from the beam path for the profile characterizations (Fig. 2 (a) and (b)) and imaging (Fig. 3). We advise building the laser combiner without the commercial speckle-reducer in a first step and evaluate then the need for further speckle reduction.

The agitation motor used here does not have ball bearings and can therefore deteriorate rapidly. We describe on the Github repository an alternative solution based on a brushless motor (RC A2212 2200KV). Using such a motor will substantially increase the durability of the agitation system and avoid regular motor exchanges.

## 4. Conclusion

In conclusion, the laser engine can deliver high-power multicolor and speckle-free illumination at an order of magnitude lower price compared to scientific-grade laser combiners. To facilitate the reproduction of such a laser engine, we made all blueprints, designs and description of the diodes, diode drivers, motor, electronics, optical and mechanical components available on Github (https://github.com/ries-lab/LaserEngine).

## 5. Materials and methods

### 5.1. Microscope

All microscopy images were acquired with a custom-built microscope. After the laser engine, an achromatic lens (10 mm, AC080-010-A) forms the image of the fiber output in the object plane of another achromatic lens (150 mm, AC254-150-A). The collimated beam is subsequently reflected on a 4x dichroic mirror (F73-410, AHF) and focused by the objective (100x, 0.49 NA, UAPON 100xOTIRF, Olympus) directly into the sample. The fluorescence dichroic and the objective, along with the stages, are housed in a custom microscope body. The fluorescence signal collected by the objective is transmitted by the 4x dichroic mirror and reflected on a mirror within the microscope body. The tube lens (U-TLU, Olympus) forms an image of the sample outside the microscope body in the object focal plane of an achromatic lens (AC254-200-A, Thorlabs). After the lens, a filter wheel allows selecting the channel (we used a band-pass 676/37, AHF). Finally, an achromatic lens (AC254-250-A, Thorlabs) focuses the light onto one half of an EMCCD camera (iXon Ultra 897, Andor). An additional 4x notch filter is placed on the camera (F40-072, AHF). In order to prevent axial drift, a focus maintaining system is implemented using a near infrared laser (785 nm, iBeam Smart, Toptica) reflected in total-internal reflection at the coverslip and detected by a quadrant photo-diode (SD197-23-21-041, Laser Components). A feedback between the photo-diode electronics and the objective stage (P-726, PI) maintains the position of the laser beam on the photo-diode fixed by moving the stage to compensate for drift. The microscope is interfaced with Micro-Manager [28] using a custom plugin, including automated activation for localization microscopy. Finally, the laser diode electronics is controlled by a FPGA (Mojo, Alchitry) compatible with Micro-Manager.

### 5.2. Power and speckle measurements

All power measurements were carried out using a commercial power meter (PM100D, Thorlabs). Coupling efficiencies reported in this manuscript were calculated by measuring the power directly at the laser outputs and after the fiber. Power densities in the sample were estimated by multiplying the light intensity measured before the objective with the objective transmittance at the specific wavelength (given in the datasheet) and dividing by the illuminated area. The latter was measured using the calibrated microscope pixel size and the image of a thin layer of dyes between two coverslips (as in [14]). The speckle contrasts were calculated by cropping the inner part of the fiber outputs (image I, 68 × 68 pixels for the round fiber, 108 × 108 pixels^2^ for the square fiber), applying a Gaussian filter with standard deviation 8 pixels to a copy of I (image G) and taking the ratio between the standard deviation of I-G and the mean <G>. A 4-pixel wide horizontal line was drawn on each fiber output image in order to produce a lateral profile in Fiji [29].

### 5.3. Sample preparation and imaging

To image the nuclear pores, we used a genome-edited U2OS cell line with Nup96-SNAP, as in [24]. The cells were prefixed for 30 seconds in 2.4% (w/v) formaldehyde in PBS (fixation buffer) on a coverslip. They were then permeabilized in 0.4% (v/v) Triton X-100 in PBS and finally incubated for 30 min in the fixation buffer again. The fixation was quenched in 100 mM NH4Cl in PBS for 5 minutes, before two washing steps in PBS of 5 minutes each. Before staining, the sample was incubated with Image-iT FX Signal Enhancer (ThermoFisher Scientific)
for 30 minutes. Staining was performed in a SNAP dye buffer (1 μM BG-AF647 (New England Biolabs; #S9136S), 1 μM DTT, and 0.5% (w/v) BSA in PBS) for 2 hours at room temperature. Finally, the samples were washed three times with PBS (5 minutes). The samples were imaged in a buffer composed of 50 mM Tris/HCL pH 8, 10 mM NaCl, 10% (w/v) D-Glucose, 500 μg/mL Glucose oxidase, 40 μg/mL Glucose catalase and 143 mM BME in H2O [30].

### 5.4. Analyses of SMLM data

All localization microscopy fitting, post-processing and rendering have been performed with SMAP. SMAP is a Matlab analysis software, freely available on Github (jries/SMAP). In particular, fitting was performed using an experimental point-spread function following [31]. After fitting, the localizations were filtered by localization precision (15 nm cut-off), estimated PSF size (150 nm cut-off) and frames (frames between 1000 and 4000). The overview reconstruction was generated using Gaussian rendered localizations with a minimum sigma of 1 nm. The zoom-in images were displayed with a minimum sigma of 6 nm. The statistics (localization precision) were computed using SMAP plugins. The average diameter of 211 nuclear pores was obtained by automatically detecting nuclear pores within 220 × 220 nm^2^ regions, filtering regions containing multiple pores or pores with insufficient number of localizations, and fitting a ring to the localizations that belong to a specific nuclear pore using least-square optimization. These steps are all available within SMAP plugins and are described in detail in [24].

## Acknowledgments

We thank Maurice Kahnwald for the sample preparation and help in imaging the biological sample and Christian Kieser (EMBL electronic workshop) for his help in building the laser diode electronics.

---

**Appendix Fig. 1.**
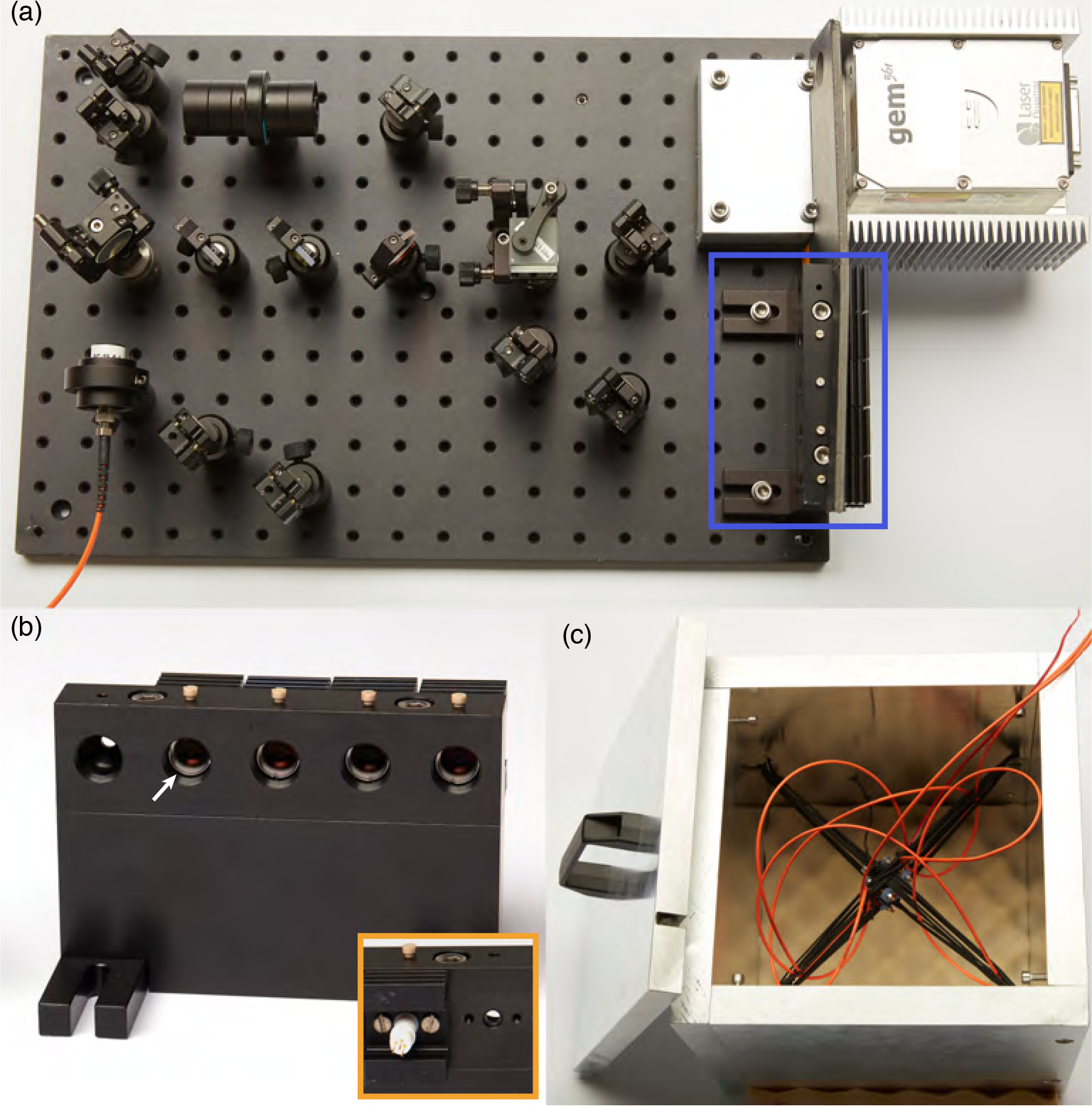
Pictures. (a) Top view of the laser engine without enclosure. The blue line indicates the position of the laser diode mount. (b) Front view of the laser diode mount with an arrow pointing at the aspherical lens. Inset picture: close-up of the laser diode mount’s back. (c) Agitation module.

**Appendix Fig. 2.**
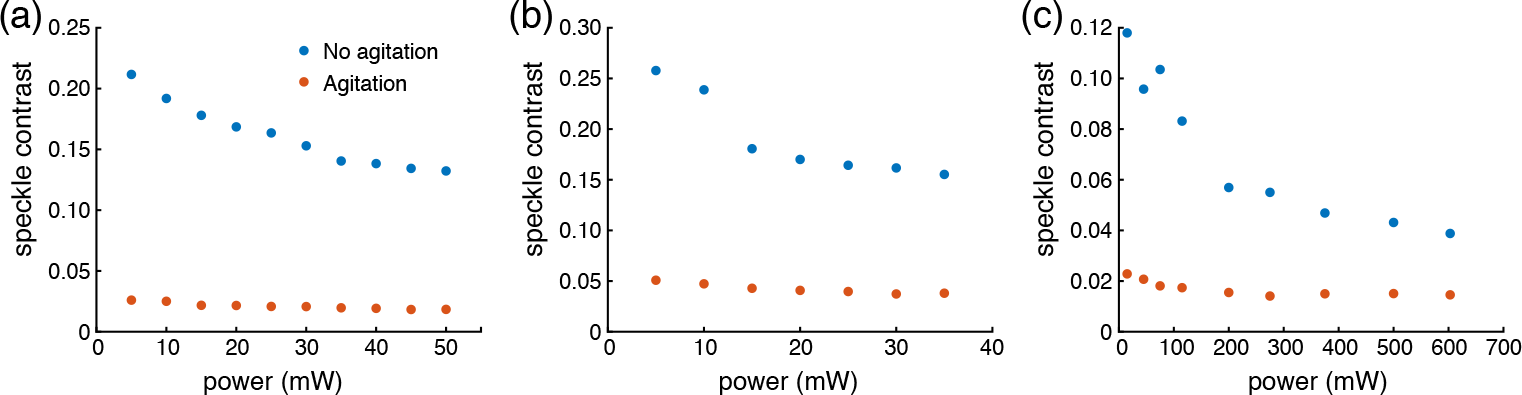
Speckle dependence on the laser diode output power. (a) Speckle contrast of the square multimode fiber at different power setpoints for the 405 nm laser diode with (red dots) and without (blue dots) agitation. The agitated data points were acquired with 5 ms exposure time. (b) Same as (a), albeit for the 488 nm laser diode. (c) Same as (a), albeit for one of the 638 nm laser diodes. All optical powers were measured directly after the laser diode mount.

**Appendix Fig. 3.**
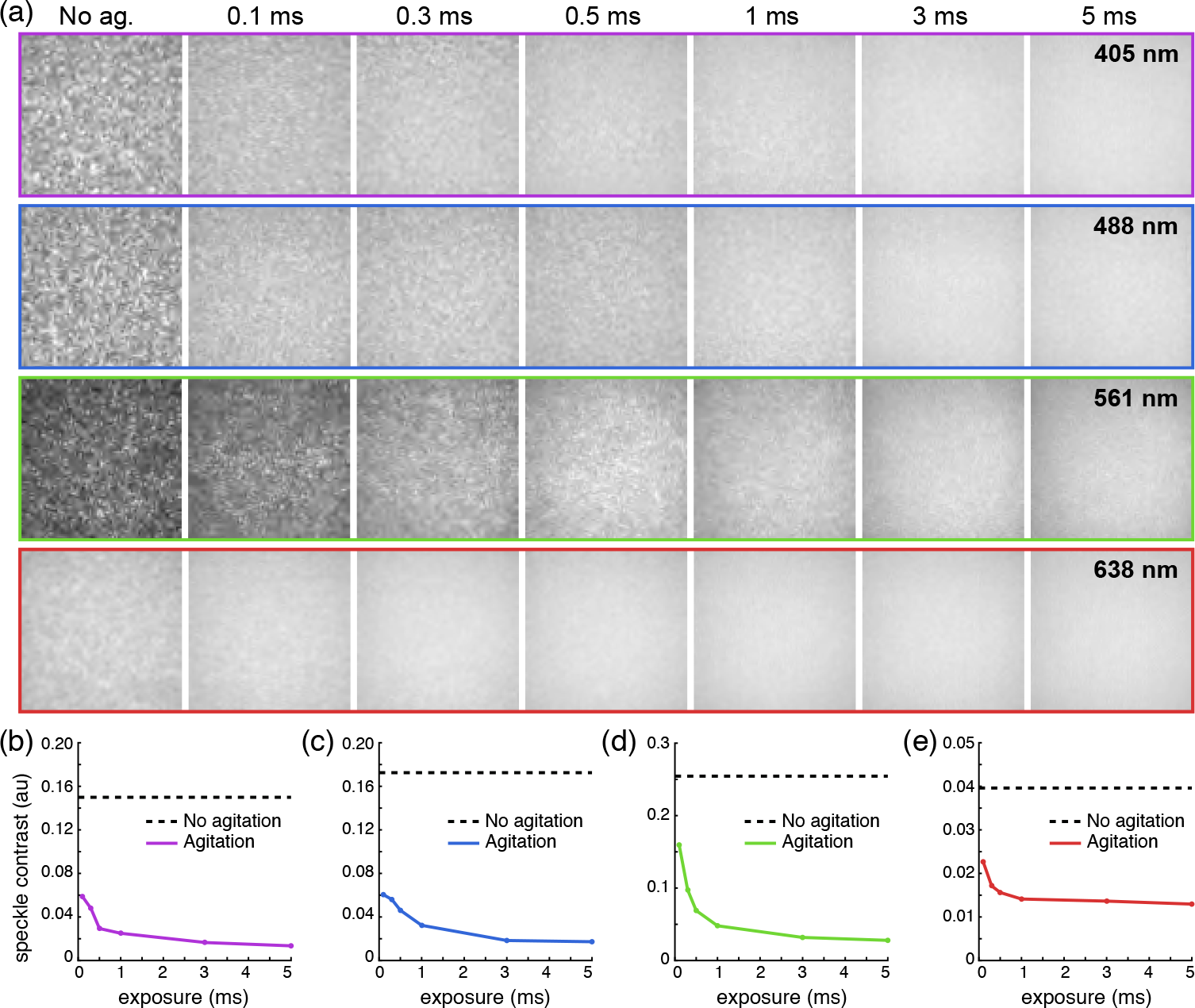
Speckle-reduction at different exposure times. (a) Central area of the square multimode fiber at different exposure times for each laser line. The leftmost column is the profile at 5 ms exposure time without agitation. The other columns are the agitated profiles at the exposure time indicated on top of each column. (b) The speckle contrast with agitation (solid line) for the 405 nm laser diode plotted against the exposure time and compared with the speckle contrast in the absence of agitation (dashed line). The output power was 50 mW. (c) Same for the 488 nm laser diode, with 30 mW laser power. (d) Same for the 561 nm dpss laser, with 200 mW laser power. (e) Same for one 638 nm laser diode, with above 400 mW of laser power.

**Appendix Fig. 4.**
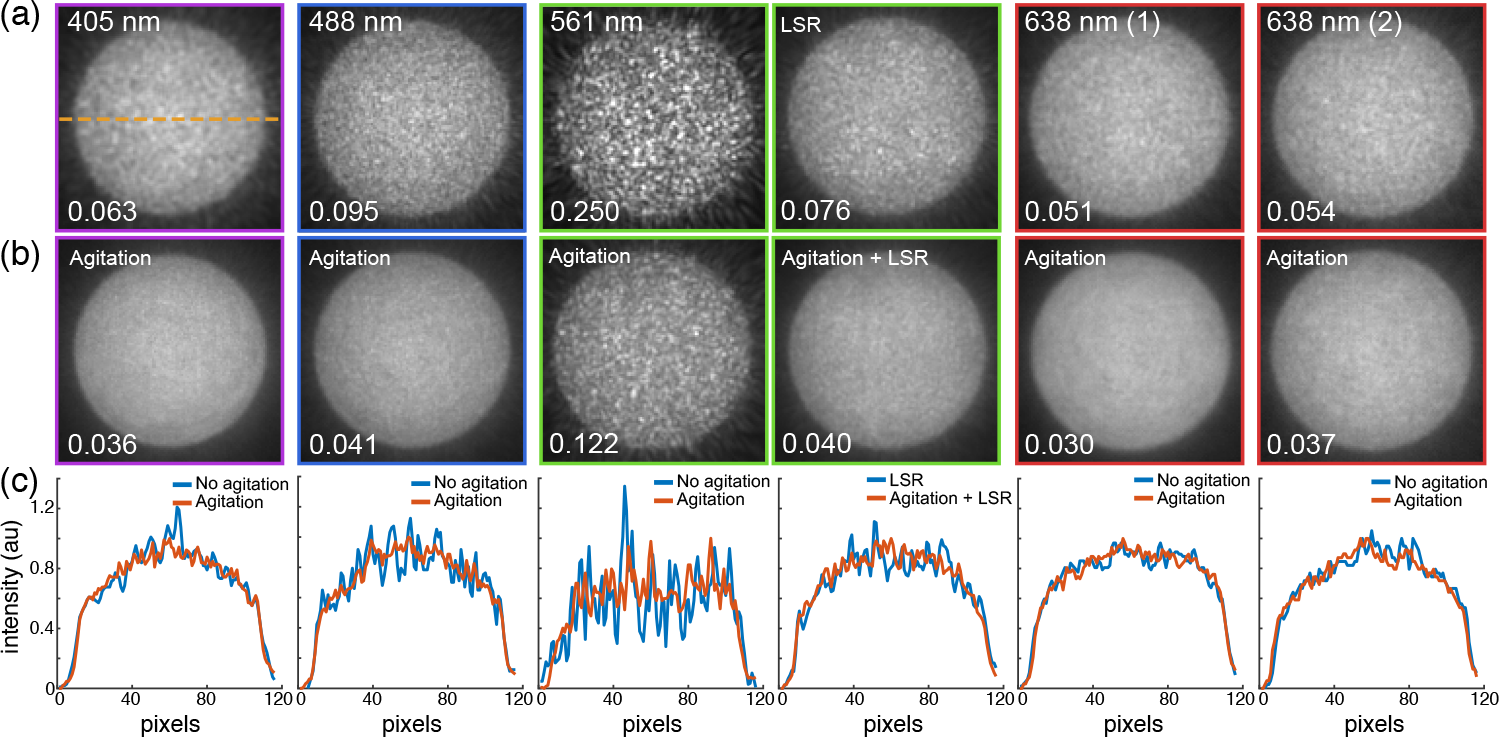
Speckle-reduction of the round multimode fiber. (a) Profiles of the round multimode fiber output without speckle reduction for the four laser lines and additionally for the 561 nm dpss laser with the LSR turned on. The speckle contrast is indicated at the bottom of each image. (b) Same profiles with agitation. (c) Lateral intensity profiles of (a) (blue curve) and (b) (orange curve) along the dashed line of (a), averaged over a width of 4 pixels. All images were acquired with 5 ms exposure time.

**Appendix Fig. 5.**
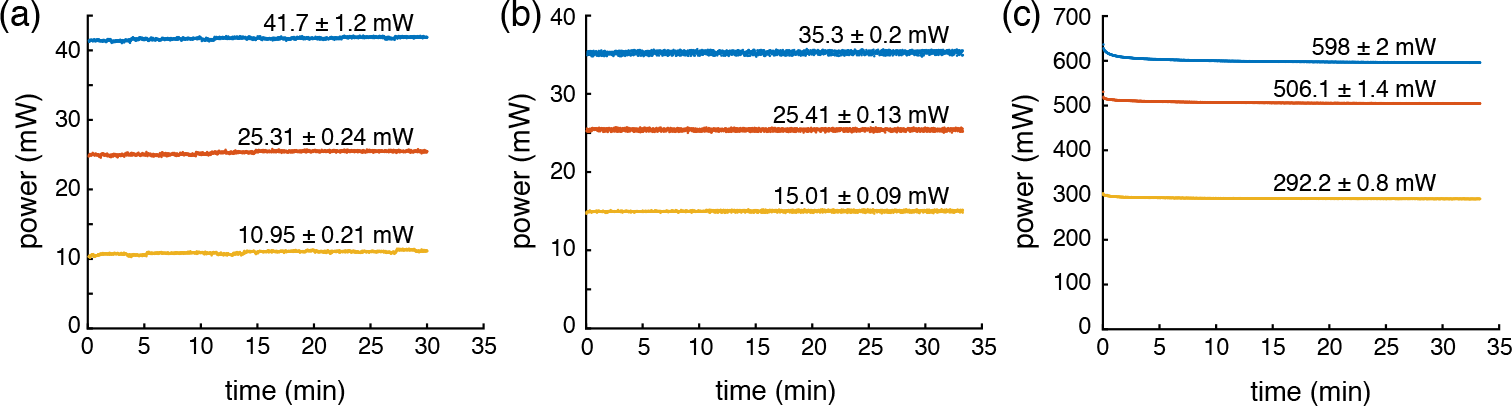
Stability of the laser diode outputs. (a) Optical power output of the 405 nm laser diode over time (sampled every second) at three different current setpoints (blue, orange and yellow curves). The mean ± the standard deviation of the last 30 min is indicated above each curve. (b) Same as (a), albeit for the 488 nm laser diode. (c) Same as (a), albeit for one of the 638 nm laser diodes.

